# Divergence in Thermostability of Arabidopsis Mitochondrial Nucleotide Exchange Factors Encoded by Duplicate Genes, *MGE1* and *MGE2*

**DOI:** 10.1101/553503

**Authors:** Zih-teng Chen, Meng-Ju Hung, Shih-Jiun Yu, Tai-Yan Liao, Yao-Pin Lin, Rita P.-Y. Chen, Chien-Chih Yang, Yee-yung Charng

**Affiliations:** Department of Biochemical Sciences and Technology, National Taiwan University, Taipei 10617, Taiwan, ROC; Agricultural Biotechnology Research Center, Taipei 11529, Taiwan, ROC; Institute of Biological Chemistry, Academia Sinica, Taipei 11529, Taiwan, ROC; Institute of Biochemical Sciences, National Taiwan University, Taipei 10617, Taiwan, ROC

## Abstract

The divergence of duplicate genes links to organismic adaptation. In *Arabidopsis thaliana* two nuclear genes encode mitochondrial GrpEs, MGE1 and MGE2, the nucleotide exchange factors of DnaK/HSP70 chaperone. *MGE1* and *MGE2* are duplicate genes originated from a whole genome duplication event. They respond differentially to high temperature; *MGE2* is heat-inducible and is required for Arabidopsis seedlings to tolerate prolonged heat stress, while *MGE1* is constitutively expressed. Heterologous expression of MGE2 but not MGE1 restored the growth of *E. coli grpE* mutant cells at elevated temperatures, suggesting that MGE2 is more thermostable than MGE1. In this study, we directly compared the thermostability of the purified recombinant MGE1 and MGE2 by circular dichroism spectroscopy. The temperature midpoints of the unfolding transition (T_m_) of MGE1 and MGE2 were about 38 and 46 °C, respectively, indicating that MGE2 is remarkably more stable than MGE1 at higher temperature. Domain swapping between the two homologous proteins showed that the N-terminal region, including an unstructured sequence and a long α-helix domain, is the major determinant of the thermostability. Although MGE2 contains a conserved sequence derived from an exonized intron within the N-terminus unstructured region, deletion of this sequence did not substantially affect protein thermostability *in vitro* and complementation of *E. coli* and Arabidopsis heat sensitive mutants. Taken together, our results suggest that Arabidopsis *MGE1* and *MGE2* had diverged not only in transcriptional response but also in the thermostability of the encoded proteins, which may contribute to adaptation of plants to higher temperatures.

## INTRODUCTION

Gene duplication provides evolutionary niches for organismic adaptation to environmental changes (Ames et al. 2010; Kondrashov 2012; Qian and Zhang 2014). Duplicate genes generated from diverse mechanisms are subjected to selection pressure that forces them to adopt different or redundant roles in host fitness. Understanding how the genes had molecularly diverged in relation to their biological functions not only sheds light on the evolution of the duplicate genes but may also on the underlying mechanism of adaptation.

The evolution of duplicate genes is closely associated with plant adaptation to environmental stresses (Panchy et al. 2016). However, the evidence pointing to this relation remains scarce. Heat stress (HS) is one of the primary environmental factors in restricting growth and development of plants, which has raised concern due to the global warming trend. In the natural environment, ambient temperature can rise swiftly with varied severity and frequency, resulting in diverse HS regimes that threaten the well-being of plants. To cope with different HS regimes, plants have evolved an array of genetic components for counteracting thermotolerance responses, *i.e*. basal thermotolerance, short- and long-term acquired thermotolerance, and thermotolerance to moderately high temperature (TMHT) (Yeh et al. 2012). Currently, we are at the infant stage in appreciating the role of thermotolerance diversity in the nature environment and the input from evolution of gene duplicates.

Among the components required for organismal thermotolerance, the best-known are the heat shock proteins (HSPs) that function as molecular chaperones or unfoldases in maintaining protein homeostasis, or proteostasis, within the cells (Bukau et al. 2006; Finka et al. 2016). Highly conserved HSPs, such as HSP70s and HSP90s, require the interaction with corresponding co-chaperones to fulfill their tasks. In plants, HSPs and their co-chaperones are often encoded by multigene families (Agarwal et al. 2001; Hill and Hemmingsen 2001; Krishna and Gloor 2001; Lin et al. 2001; Miernyk 2001). HSP70s play a critical role in the proteostasis networks in various cellular compartments of the eukaryotes (Hartl et al. 2011). They require the sequential interaction with two types of co-chaperones: J-domain proteins that stimulate the ATPase activity of HSP70 and nucleotide exchange factors (NEFs) that facilitate the replacement of the bound ADP by ATP (Mayer and Bukau 2005). The conversion between the ADP-bound and ATP-bound states changes the affinity of HSP70 to its polypeptide substrates, which promotes protein folding.

In the *A. thaliana* nuclear genome, there are 14 *HSP70* genes encoding chaperone proteins targeted to cytosol, ER lumen, mitochondrion matrix, and plastid stroma (Lin et al. 2001; Sung et al. 2001). The mitochondrial and the plastid HSP70s are derived from eubacterial origins and are classified as the DnaK type by having the GrpE signature loop within the N-terminal nucleotide binding domain (Brehmer et al. 2001; Lin et al. 2001). GrpE is the NEF specific for the interaction with the DnaK-type HSP70. The GrpE-associated HSP70 machineries are involved in the folding of the nascent protein peptides, organellar protein translocation, and disaggregation and refolding of the denatured proteins (Mayer and Bukau 2005).

Previous studies on the two *A. thaliana* mitochondrial GrpE proteins, MGE1 and MGE2, showed that they are encoded by duplicate genes generated from a whole-genome duplication event around the K-T boundary (Hu et al. 2012). Interestingly, the duplication and retainment of two *MGE* genes also occurred independently in many plant species, suggesting a common evolutionary tendency of the genes (Hu et al. 2012). In Arabidopsis and tomato, *MGE1* is constitutively expressed, while *MGE2* is heat-induced, suggesting subfunctionalization of the *MGE* genes by differential transcription. In Arabidopsis, expression of *MGE2* is dependent on HSFA1s, the master transcription regulators of HS response, while expression of *MGE1* is independent of HSFA1s (Liu et al. 2011; Hu et al. 2012). *MGE2* is dispensable for normal growth and development but specifically required for TMHT in Arabidopsis. Arabidopsis MGE2 contains a motif enriched in Lys, Arg, and Ser residues (KRS-enriched motif) coded by a nucleotide sequence originated from an exonization event, in which intron 2 becomes part of an exon (Fig. 1). Intriguingly, in other higher plant species, MGE2 isoforms containing the conserved KRS-enriched motif can also be produced by alternative splicing of the corresponding intron (Fig. 1 and (Hu et al. 2012)). However, the role of the KRS-enriched motif is unclear. Heterologous expression of Arabidopsis MGE2 but not MGE1 complemented the heat sensitive phenotype of an *E. coli* mutant (DA16) with a loss-of-function *GrpE* gene (Hu et al. 2012). These observations suggest that *MGE1* and *MGE2* diverged not only at transcriptional regulation but also at the protein level. One possible direction of divergence may result in difference in protein thermostability of the MGEs. We hypothesized that MGE2 has a higher thermostability than that of MGE1.

**Fig. 1.**
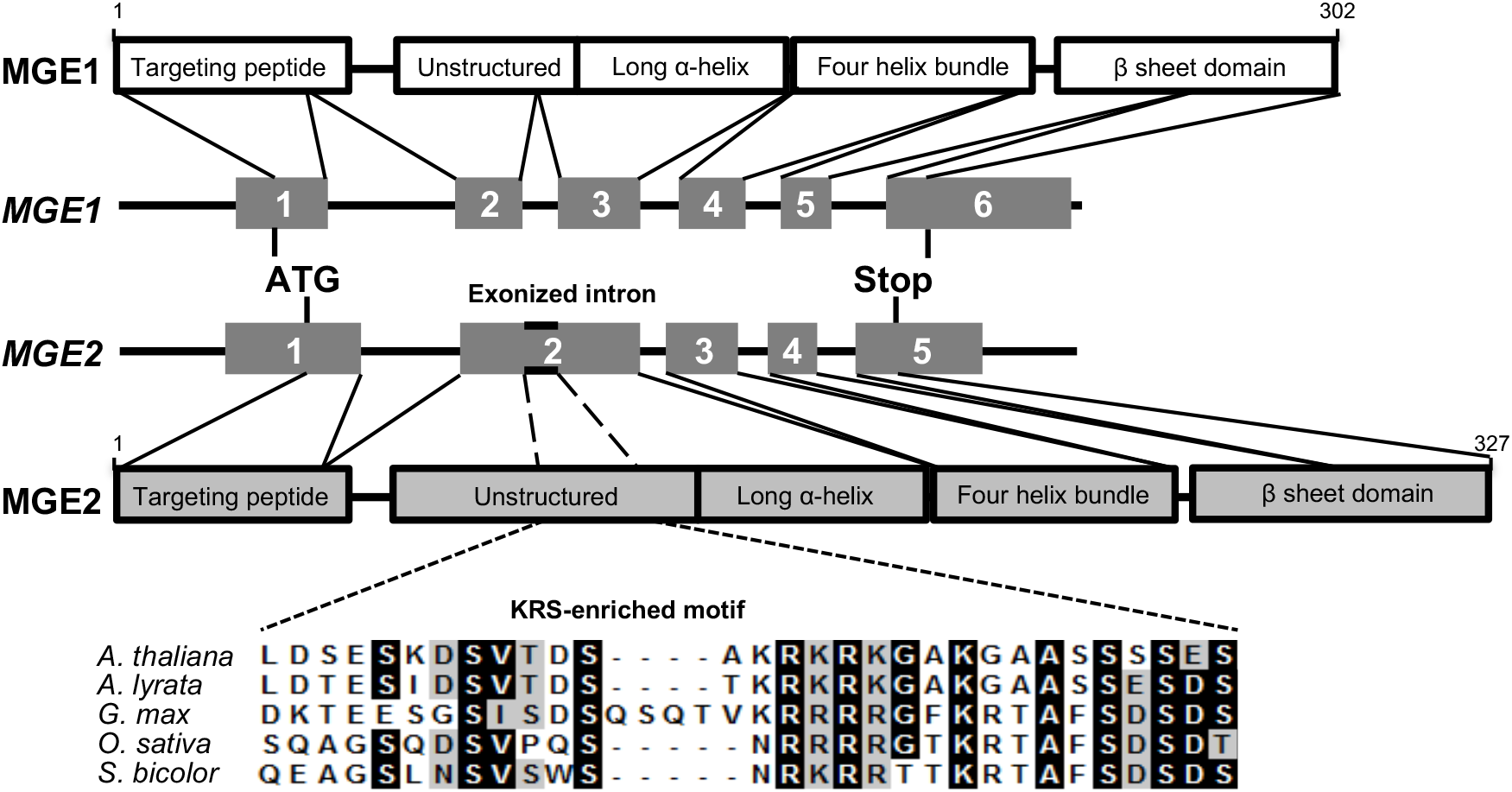
Schematic structures of Arabidopsis *MGE* genes and corresponding protein features. The exons of the Arabidopsis genes are numbered in dark-gray boxes, and the corresponding protein structures are indicated. The KRS-enriched motifs encoded by exonized or alternatively spliced intron in the *MGE2* genes from different species (*Arabidopsis thaliana, Arabidopsis lyrate, Glycine max, Oryza sativa*, and *Sorghum bicolor*) are aligned and shown below. The identical and similar amino acid residues are marked in black and gray backgrounds, respectively. The sources of the sequences were described in Hu et al. (2012).

To test our hypothesis, we compared the thermostability of the recombinant MGE1 and MGE2 proteins purified for circular dichroism (CD) spectroscopy analysis. Swapping of the three major domains between the two MGEs was performed to identify the features that affect protein thermostability. We also examined the *in vitro* and *in vivo* effect of deleting the KRS-enriched motif of MGE2. Our results showed that MGE2 has a substantially higher thermostability than MGE1, mainly contributed by the N-terminal portion including the unstructured sequence and the long a-helix domain, suggesting divergence of the MGE proteins in thermostability. However, the conserved KRS-enriched motif located within the N-terminal unstructured sequence does not play a substantial role. Overall, this study provides a better insight on the evolution of the duplicated *MGE* genes in Arabidopsis. The divergence of protein thermostability in MGEs may facilitate adaptation of plants to prolonged heat stress.

## RESULTS

### Functional confirmation and purification of *Strep*-tagged MGEs

To facilitate the purification of proteins for CD analysis, the recombinant MGE1 and MGE2 were engineered to fuse to an eight-residue affinity tag, *Strep-tag* II (WSHPQFEK) at the C-terminus (Schmidt and Skerra 2007). To examine whether the *Strep-tag* affected MGE function, complementation assay using the *E. coli grpE* mutant, DA16, was performed. DA16 carries a GrpE (G122D) mutation and exhibited a heat-sensitive phenotype (Grimshaw et al. 2005; Harrison et al. 1997). Transformation of DA16 with the constructs encoding mature MGE1 and MGE2 rescued the heat sensitive phenotype to differential extent (Supplemental Fig. 1), which is consistent with previous report (Hu et al. 2012). Expression of the *Strep*-tagged MGE proteins in D16 conferred thermotolerance similar to the levels conferred by the recombinant proteins without the affinity tag (Supplemental Fig. 1), suggesting that the *Strep*-tag does not interfere with the function of MGEs. The *Strep*-tagged MGEs were then purified from the crude extract of *E. coli* cells expressing the recombinant proteins by affinity chromatography using a *Strep*-trap column. The *Strep*-tagged MGEs can be purified as the predominant protein after the affinity column elution. However, minor proteins were copurified after this step, which were removed by ion exchange and gel filtration (Supplemental Fig. 2). The purity of the recombinant proteins were estimated to be > 90% by SDS-PAGE and Coomassie Blue staining.

### Recombinant MGE2 is more thermal stable than MGE1 *in vitro*

To compare the thermostability of the two recombinant MGE proteins, we used CD spectroscopy to determine the denaturation tendency of the purified proteins in response to increasing temperature. Changes of secondary structures were monitored under temperatures varied from 10 °C to 90 °C for thermo-induced protein unfolding, and then from 90 °C to 10 °C for protein refolding. In general, proteins with high percentage of α-helices like MGEs present two negative ellipticity at 208 and 222 nm along with one maximum signal at 192 nm in CD spectra. Therefore comparing the CD signal at 222 nm is commonly used as an indicator of the change in the α-helical structure under treatments. As shown in Fig. 2A, the thermal transition of MGE1 began at about 28 °C. As the temperature increased, the MGE1 protein gradually lost its α-helical structure. After the temperature reached 50 °C, folding of the α-helical structure of MGE1 was no longer observed. The CD spectra indicate that the unfolding of MGE1 has a transition midpoint (T_m_) at approximately 38.2 °C (Fig. 2A). Moreover, the CD spectra during the refolding process by cooling the unfolded MGE1 from 90 °C to 10 °C had lower amplitude than the original one, indicating part of the protein did not regain the α-helical conformation and might aggregate after thermal unfolding. On the other hand, unfolding of MGE2 began and completed at higher temperatures (about 40 °C and 60 °C, respectively), and the T_m_ of MGE2 was about 46.2 °C (Fig. 2C). Of note, most of the α-helical structure of MGE2 protein was regained after cooling to 10 °C. Our results demonstrate that MGE2 is more stable than MGE1 at elevated temperature *in vitro*. These results also suggest that the unfolding of MGE2 is reversible, whereas unfolding of MGE1 is partially irreversible when heated to 90 °C.

**Fig. 2.**
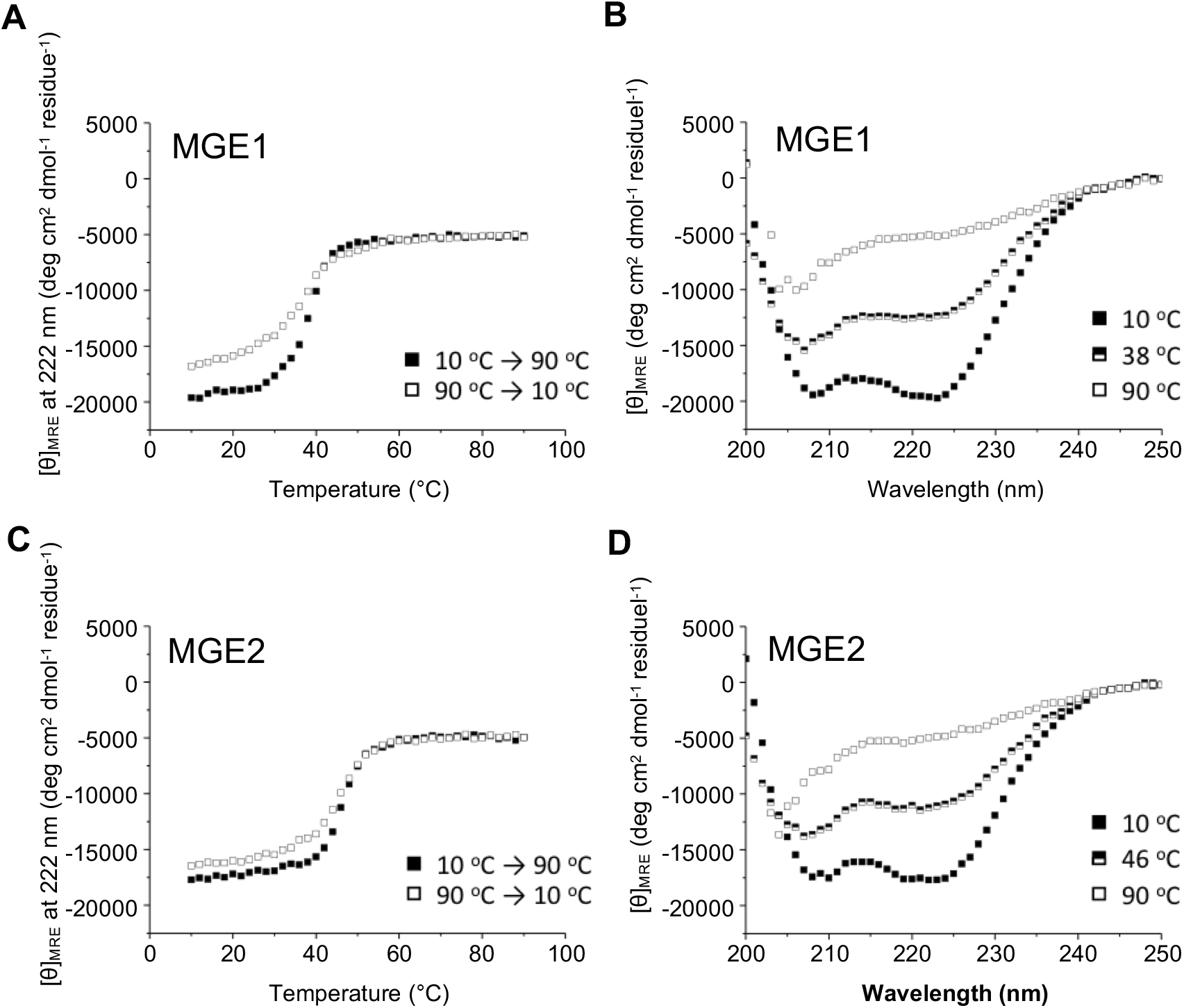
The thermal unfolding/refolding transition of the purified recombinant MGE1 and MGE2. Changes of the α-helical structure of MGE1 (A) and MGE2 (C) in CD spectra were monitored at 222 nm and expressed in mean residue ellipticity (MRE). MGE1 (233 ng/μL) and MGE2 (200 ng/μL) samples were heated from 10 °C to 90 °C (black square) in a cuvette with a path length of 1 mm, then cooled from 90 °C to 10 °C (white square). CD spectra of purified MGE1 (B) and MGE2 (D) proteins were measured at 10 °C (solid black square), 90 °C (empty white square) and the midpoint of thermal transition (half-black square).

### Domain swapping revealed the predominant feature for thermostability of MGEs

Based on the sequence homology shared with *E. coli* GrpE, whose crystal structure is available (Harrison et al. 1997), Arabidopsis MGE1 and MGE2 are predicted to consist from N-to C-terminus an unstructured sequence, a long α-helix domain, a four-helix bundle domain, and a β-sheet domain (Fig. 1). To determine which feature accounts for the difference in thermostability of MGE1 and MGE2, we swapped the unstructured sequence plus the long α-helix domain, the four-helix bundle, and the β-sheet domain between MGE1 and MGE2 to generate six chimeric proteins as indicated in Fig. 3. The chimeric MGE proteins were purified and analyzed by CD spectroscopy (Supplemental Fig. 3). Our results showed that the chimeric proteins contained the N-terminal unstructured sequence plus the long α-helix domain derived from MGE2 have higher T_m_ in general (Fig. 3), indicating that this region is the predominant feature for the thermostability *in vitro*.

**Fig. 3.**
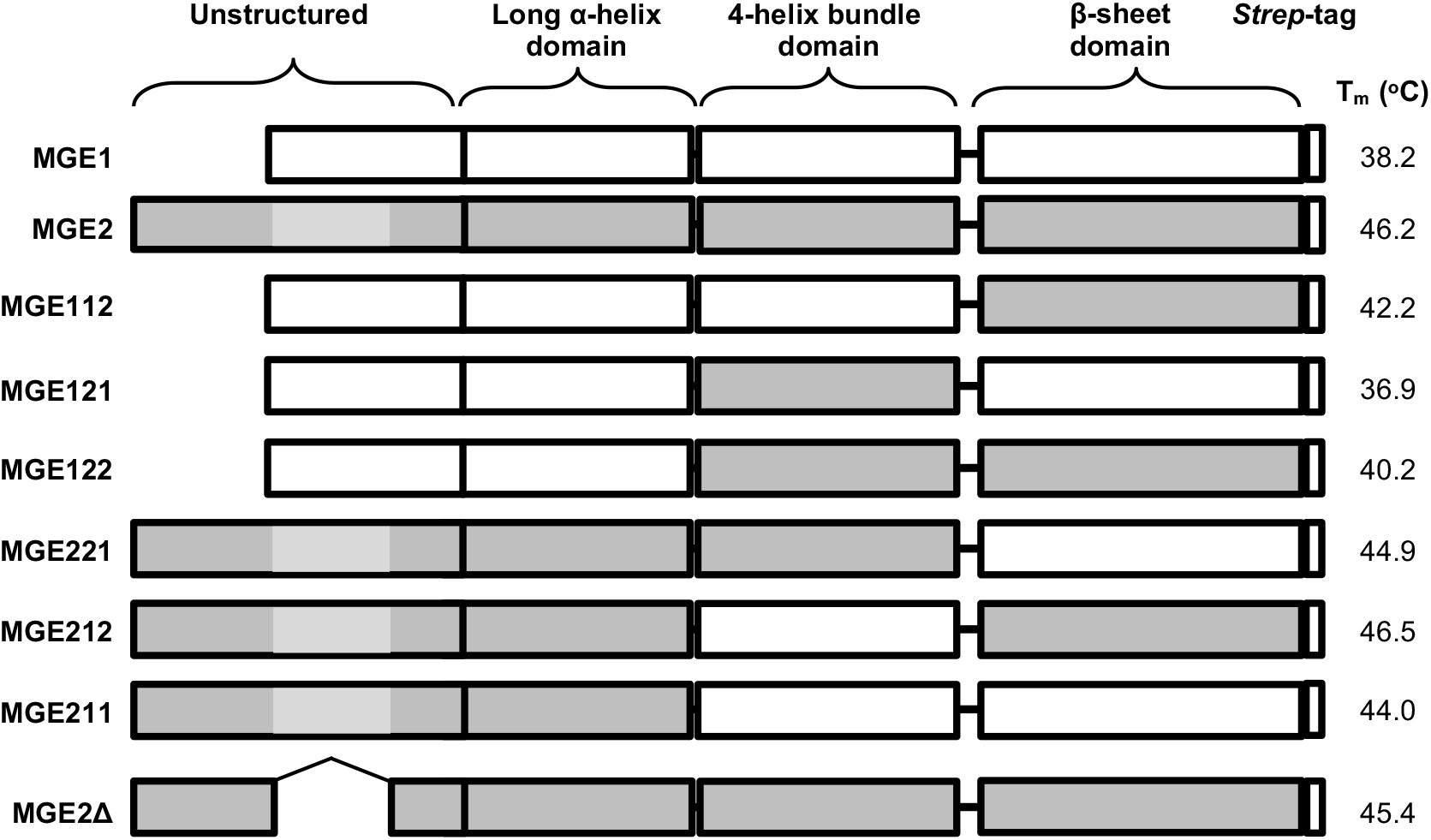
Schematic structures of domain-swapped and truncated MGE proteins and respective T_m_ values. The domains of the *Strep*-tagged MGE1 and MGE2 proteins were shown in white and dark gray rectangles, respectively. Three major regions were swapped between MGE1 and MGE2, *i.e*. the unstructured sequence plus the long α-helix domain, the four-helix bundle, and the β-sheet domain, to generate six chimeric proteins denoted as MGE112, 121, 122, 221, 212, and 211. MGE2Δ is a truncated MGE2 without the KRS-enriched motif (shown in light gray rectangles).

To see whether these chimeric proteins also show similar thermal response *in vivo*, the plasmids expressing the chimeric MGEs proteins were transformed into *E. coli* DA16 for complement assay. Consistent with the *in vitro* results, *E. coli* DA16 expressing the chimeric proteins with the N-terminal unstructured sequence plus the long α-helix domain of MGE2 could grow nicely at 43 °C, while the chimeric proteins with the features from MGE1 did not confer tolerance to the high temperature condition (Fig. 4A).

**Fig. 4.**
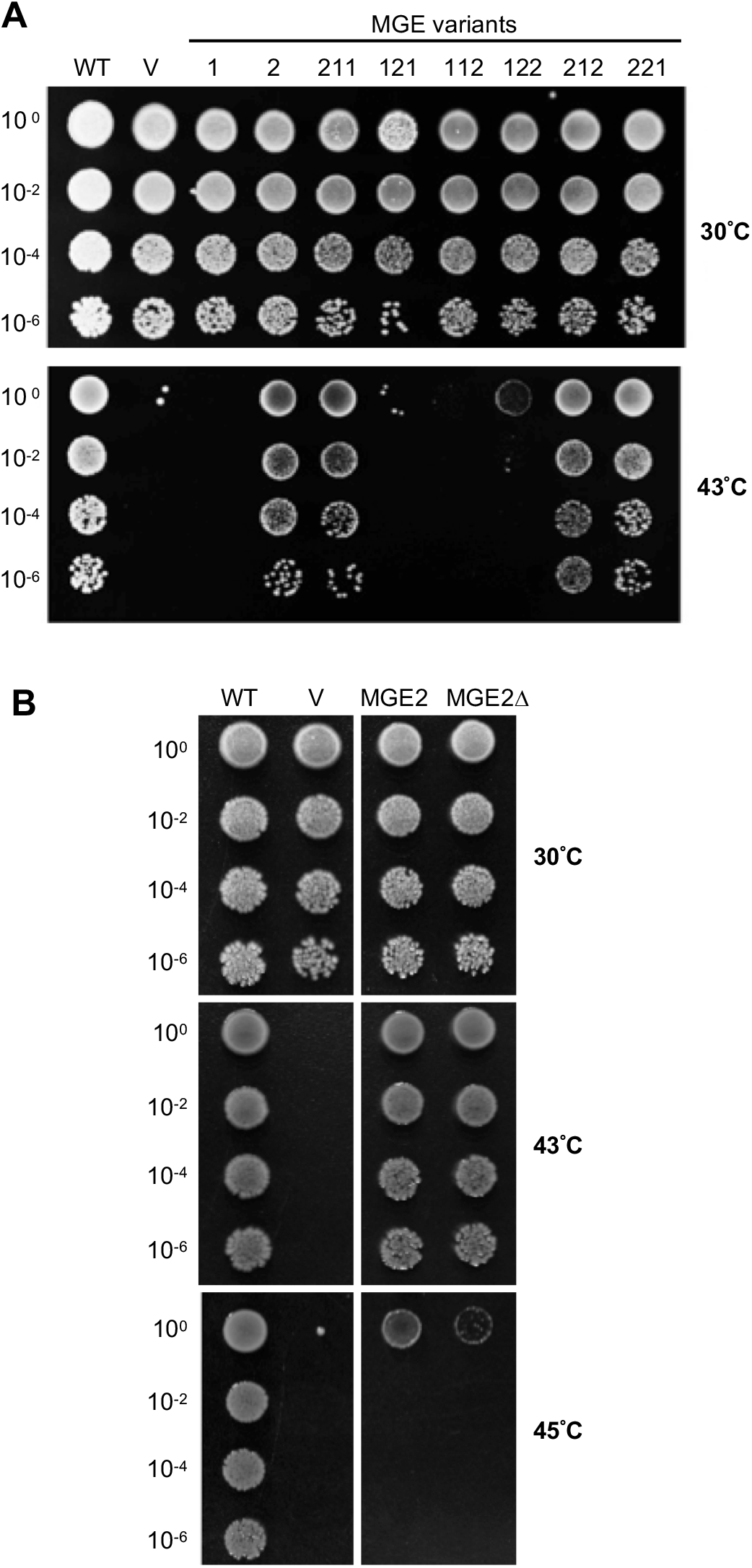
Thermotolerance assay of the *E. coli grpE* mutant (DA16) complemented with MGE variants. Complementation was assessed by the growth of *E. coli* cells at indicated temperatures. Cell cultures were serially diluted by 100-fold and spotted onto LB plates containing ampicillin for overnight. A, DA16 cell lines transformed with empty vector, *Strep*-tagged MGE1 or MGE2 construct were labeled as V, 1, or 2 respectively. The wild-type strain DA15 (WT) with an isogenic background to DA16 was also transformed with the same empty vector as a control. Cell lines transformed with the constructs encoding domain-swapped MGEs were indicated as illustrated in Fig. 4. B, The growth of DA16 cell lines transformed with the constructs encoding MGE2 and MGE2Δ was compared with that of WT and DA16 with vector control (V).

One obvious difference between the N-terminal unstructured sequence of MGE1 and MGE2 is the presence of the intron-derived KRS-enriched motif in the latter (Fig. 1). To elucidate the function of the KRS-enriched motif, a shorten version of MGE2, designated as MGE2Δ, was generated by deleting this peptide sequence of 30 amino acid residues. The T_m_ of MGE2Δ protein was about 45.4 °C (Fig. 3), which was close to the T_m_ of MGE2 (46.2 °C). This result suggests that the KRS-enriched motif does not contribute substantially, if any, to the difference in thermostability between MGE1 and MGE2.

### MGE2 without the KRS-enriched motif remains functional in conferring thermotolerance in *E. coli* and Arabidopsis

Since the KRS-enriched motif is highly conserved in plant MGE2s, we speculated that it may be required for organismic thermotolerance. Hence, complementation experiments in *E. coli* DA16 and Arabidopsis *mge2* were performed. Fig. 4B shows that MGE2Δ could complement the growth of *E. coli* DA16 at high temperatures indistinguishable from the wild type MGE2, suggesting that the peptide derived from the in-frame intron in Arabidopsis MGE2 does not play a critical role in its function associated with thermotolerance in *E. coli* cells. Similarly, MGE2Δ could restore the TMHT of Arabidopsis *mge2* as well as the wild type protein (Fig. 5A). The immunoblot showed a faster migration of MGE2Δ on SDS-PAGE than that of MGE2 expressed in the transgenic lines, indicating a size reduction by deletion of the KRS-enriched motif protein (Fig. 5B). The level of MGE2Δ was accumulated in response to prolonged HS like the wild-type, suggesting that the peptide sequence is also not essential for protein stability under stress condition.

**Fig. 5.**
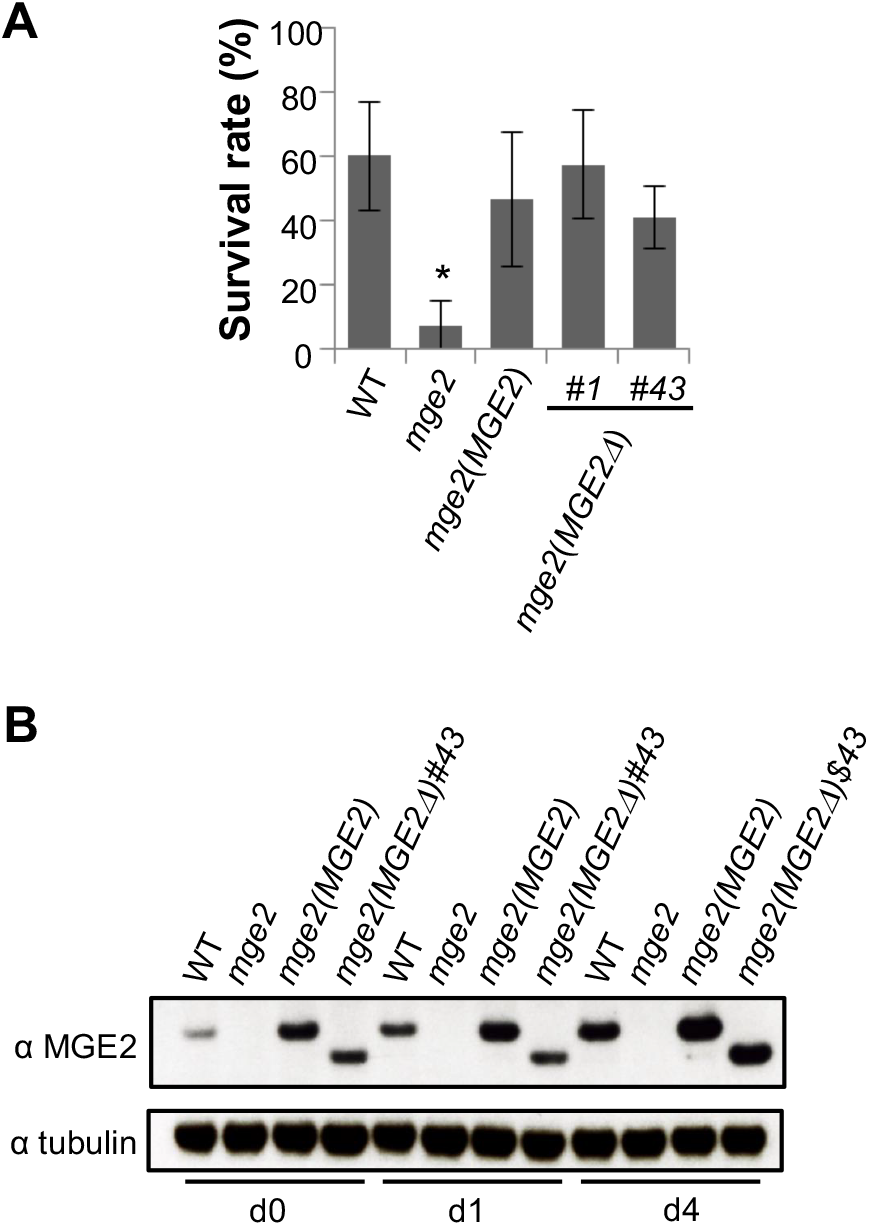
Complementation analysis of *Arabidopsis mge2* knockout mutant transformed with *MGE2* or *MGE2Δ*. **A**, Comparison of the survival rates between WT and the transgenic lines after TMHT assay. *mge2*(*MGE2Δ*)#1 and #43 are two independent lines. Results are the means of three replicates ± SD (n ≥ 25). *P < 0.01 (versus wild-type plants, Student’s t-test). **B**, Immunoblots of total proteins extracted from Arabidopsis seedlings harvested after heat treatment at 35 °C for 1 (d1) or 4 days (d4). The sample collected before heat treatment was denoted as d0. The transgenic lines containing the *MGE2* and *MGE2Δ* transgenes in the *mge2* background were denoted as *mge2(MGE2)* and *mge2*(*MGE2Δ*), respectively. Tubulin level was shown as a loading control.

## DISCUSSION

In this study, we compared the thermostability of recombinant Arabidopsis MGE1 and MGE2 *in vitro* and found that MGE2 was substantially more stable than MGE1 at high temperature. Although these proteins were fused to a *Strep-tag* to facilitate purification, the tag does not affect the protein thermostability considerably since no obvious difference in their capacity in conferring thermotolerance in *E. coli* with or without the tag (Supplemental Fig. 1). CD spectra showed that the configuration of MGE1 started to alter at around 28 °C, while that of MGE2 was not affected up to 40 °C (Fig. 2). This result is consistent with the fact that the heterologous expression of Arabidopsis MGE1 in *E. coli* DA16 could not support the growth of the bacterial mutant at 43 °C (Fig. 4A and (Hu et al. 2012)). At this temperature, which is higher than its T_m_ (38.2 °C), MGE1 is likely unstable *in vivo* and no longer functional as an NEF of mitochondrial HSP70. The works on *E. coli* GrpE and yeast mitochondrial GrpE, Mge1p, had shown that thermally denatured GrpE and Mge1p do not interact stably with DnaK and mitochondrial HSP70, respectively, and are unable to regulate the protein folding activity of the chaperone (Grimshaw et al. 2003; Moro and Muga 2006). In contrast, heterologous expression of Arabidopsis MGE2 could rescue the *E. coli* mutant at 43 °C (Fig. 4), which is below the T_m_ (46.2 °C) of the protein (Fig. 2C, D). These results indicate that Arabidopsis MGE2 is more thermal stable than MGE1 both *in vitro* and *in vivo*.

The thermostability of MGEs shown here also supports their functions *in planta. MGE1* is expressed in Arabidopsis at normal growth temperature when *MGE2* expression is hardly detected (Hu et al. 2012), suggesting that MGE1 plays a housekeeping role in proteostasis under normal condition. Given its low thermostability, MGE1 is unstable or less efficient at elevated temperature. Hence, the heat-inducible and more thermostable MGE2 replaces MGE1 in maintaining proteostasis under prolonged HS. Our results demonstrate that, in addition to the divergence in expression profile (Hu et al. 2012), the divergence in protein thermostability of MGE1 and MGE2 is discernible in the evolution of the *MGE* duplicates.

Given that Arabidopsis MGEs can complement the function of *E. coli* GrpE and the close similarity in amino acid sequence, MGE1 and MGE2 should also contain three structured domains: a long α-helix domain, a four-helix bundle domain, and a β-sheet domain, based on the crystal structure determined for *E. coli* GrpE (Harrison et al. 1997). The sequence similarities between MGE1 and MGE2 are about 53 % for the long α-helix, 93 % for the four-helix bundle, and 89 % for the β-sheet domain. The similarity between the N-terminus unstructured regions of MGE1 and MGE2 is even lower than 50%. Domain swapping analysis revealed that the N-terminal portion of MGE proteins, including the N-terminus unstructured region and the long α-helix domain, is the major determinant of the thermostability (Fig. 3). This result is consistent with the fact that the N-terminal portion of the MGEs share the least sequence homology among the three swapped domains. Although an obvious difference between the N-terminus unstructured regions of the proteins is the presence of the KRS-enriched motif in MGE2 (Fig. 1), we excluded the possibility that this peptide sequence is a major factor of the high thermostability of MGE2 since the motif only slightly increase the T_m_ value of MGE2 by 0.8 °C (Fig. 3), we did not observe a substantial difference in conferring TMHT by MGE2 with or without the KRS-enriched motif *in planta* (Fig. 5A). The thermotolerance assay may not be sensitive enough to reveal the effect of the peptide deletion. Alternatively, the KRS-enriched motif might have a function other than thermotolerance.

*E. coli* GrpE has been proposed to be the thermosensor of the DnaK chaperone machinery as its configuration undergoes reversible thermal transition, which regulates the NEF activity of GrpE (Gelinas et al. 2002; Grimshaw et al. 2003). Thermodynamic analysis indicated that *E. coli* GrpE possesses two well-separated thermal transitions with midpoints at ~48 °C and ~80 °C (Grimshaw et al. 2001). Unfolding of the long α-helix domain takes place during the first transition, whereas the second transition is correlated to the unfolding of the four-helix bundle domain (Gelinas et al. 2002). Hence, the long α-helix domain plays a determining role in sensing changes of ambient temperature for the bacterial protein. However, the CD spectra suggested both Arabidopsis MGE1 and MGE2 undergo a single thermal transition (Fig. 2A, C). Single thermal transition was also shown in *S. cerevisiae* Mge1p (Moro and Muga 2006). It was suggested that the unfolding of both the long α-helix domain and the four-helix bundle domain occurs simultaneously in a very narrow temperature range. Hence, it might be also the case for the plant proteins.

The duplication and retention of the *MGE* genes occurred independently in many higher plants (Hu et al. 2012). There must be a common selection pressure that forced the ancestors of the higher plant species adopting a common evolutionary trajectory of the *MGE* genes. Given the Arabidopsis *MGE* genes duplicated during the K-T boundary, when global climate change was predicted to occur, we suspected that climate change could have played a role in *MGE* evolution. Why plants kept both copies of the *MGE* genes after duplication? Our works on the Arabidopsis MGEs suggest that the *MGE* genes underwent divergence at transcriptional regulation and protein thermostability, which might be essential for plants to adapt to a wider spectrum of temperature change. Our findings also point to a possibility that manipulation of the thermostability of MGE may contribute to improving the organismic thermotolerance, which however requires further investigation.

## MATERIALS AND METHODS

### Constructs for production of *Strep*-tagged recombinant MGEs and their variants

The Arabidopsis *MGE* cDNA sequences were amplified by RT-PCR using two primer pairs, AtMge1-bs1 plus AtMge1-bs4 for *MGE1* and AtMge2-bs3 plus AtMge2-bs4 for *MGE2* (primer sequence information provided in Supplemental Table I), as previously described (Hu et al. 2012). The primers contain restriction sites for cloning into pASK3 plus vector (IBA BioTAGnology) to generate an in-frame fusion of the *MGE* cDNA and the *Strep*-tag sequence. The resulting plasmids for producing recombinant *Strep*-tagged MGE1 and MGE2 proteins in *E. coli* were named pAMS1 and pAMS2, respectively.

For generation of domain-swapped MGE proteins, the cDNA sequences corresponding to the swapped domain and the original domains were amplified separately with pAMS1 and pAMS2 as templates by using the sticky-end PCR method (Zeng 1998) or traditional PCR. The primer sequences for PCR were provided in Supplemental Table I. The PCR product of the swapped domain was designated as the “insert”, and the two original domains were PCR amplified with the rest of the expression vector marked as the “vector” fragment. The “insert” and “vector” fragments were then ligated after appropriate processing and transformed into *E. coli* DH5α cells. The plasmids for expressing the *Strep*-tagged and domain-swapped MGEs (Fig. 4) were named pAMS-211 (for MGE211), pAMS-121 (for MGE121), pAMS-112 (for MGE112), pAMS-122 (for MGE122), pAMS-212 (for MGE212) and pAMS-221 (for MGE221).

For generation of MGE2Δ, the DNA sequence corresponding to the exonized intron of *MGE2* were removed by PCR using pAMS2 as template and primers AtMGE2IntD-FP and AtMGE2IntD-RP (Supplemental Table I), then blunt-end ligation. The plasmid for expressing the *Strep*-tagged MGE2Δ protein was named pAMS2D. All the constructs were sequenced to confirm no missense or nonsense mutation.

### Purification of MGE proteins

*E. coli* BL21 (DE3) cells transformed with desired plasmids were grown in LB medium containing 100 μg/mL ampicillin at 37 °C. Protein expression was induced by adding 0.2 μg/mL of anhydrotetracycline to the culture when A600nm reached 0.6 and incubated at 37 °C for 4 h. The *E. coli* cells were pelleted by centrifugation at 4,500 xg at 4 °C for 12 min, resuspended in buffer A (20 mM sodium phosphate, 280 mM NaCl, 6 mM KCl, pH 7.4) with 1 mM EDTA and protease inhibitor (Roche cOmplete, Cocktail Tablet), and subsequently incubated with 1 mg/mL lysozyme at 4 °C for 30 min and further broken with French press (Thermo Scientific). The total lysate was centrifuged at 98,000 × g for 30 min at 4 °C. The supernatant was loaded onto a 1 mL StrepTrap HP column (GE Healthcare) and then washed with 50 mL of buffer A. The Strep-tagged MGE proteins were eluted with buffer A supplemented with 2.5 mM desthiobiotin (Novagen and Sigma-Aldrich). The protein composition of the eluent was verified by western blot using *Strep*-tag II monoclonal antibodies (Novagen). The fractions containing MGE protein were pooled and concentrated to 1 mL by using a centrifugal concentrator (Sartorius, 10 kDa MWCO, Vivaspin turbo 15). The recombinant proteins were further purified by ion exchange and gel filtration chromatography with ÄKTAprime plus system. The ion exchange column (DEAE FF, GE Healthcare) was equilibrated with 50 mM MES buffer (pH 5.8), and the absorbed proteins were eluted stepwise with the MES buffer containing 0.1, 0.2, 0.3, and 0.4 M NaCl. The protein composition of the eluent was verified by western blot using *Strep*-tag II monoclonal antibodies (Novagen). The fractions containing MGE protein were pooled and concentrated to 1 mL by using a centrifugal concentrator (Sartorius, 10 kDa MWCO, Vivaspin turbo 15). For gel filtration, the concentrated MGE proteins were loaded onto a Sephacryl S-200 High Resolution column and eluted with buffer A at flow rate 0.2 mL/min. The fractions with high A_280_ signals and abundance of MGE were pooled and concentrated. Protein concentration was determined by Bradford protein assay (Thermo Fisher Scientific). The purity of the purified MGE protein was estimated by SDS-PAGE and Coomassie Blue staining.

### Circular dichroism spectroscopy

Protein samples were diluted with buffer A to about 200 to 250 ng/μL. The secondary structure and thermostability were analyzed by Jasco J-815 CD spectrometer and using a cuvette with a 1 mm path length. The CD spectrums were determined in the range of 200-250 nm every 1 nm increment and analyzed every 2 °C ranging from 10 to 90 °C. The temperature slope of the heating process was set to 10.0 °C/min and the delay time before every measurement was 30 sec. The scanning speed was 50 nm/min and the bandwidth was set to 1 nm. The T_m_ value and the denaturation curve were analyzed by Excel and Origin Pro.

### Functional complementation of *E. coli* DA16 with Arabidopsis MGE variants

The plasmids for expressing MGE1, MGE2, and different MGE variants were transformed into *E. coli* DA16 line as previously described (Hu et al. 2012). The transformed DA16 lines were grown at 30 °C in Luria-Bertani (LB) medium containing ampicillin (100 μg/mL) to about OD_600_ 0.8. The liquid cultures were 10-fold serially diluted, and 4 μL of each dilution was dropped onto LB plate containing ampicillin. The plates were then incubated in a growth chamber at the designated temperature overnight.

### Transformation of Arabidopsis *MGE2* knockout mutant

To generate the *mge2*(*MGE2*) and *mge2*(*MGE2Δ*) transgenic lines, a 3,694 bp fragment of the *MGE2* (At4g26780) gene was amplified by PCR with a pair of primers, AtMge2 5′P-1F and AtMge2 3′P-1R (Supplemental Table I), using Arabidopsis wild-type (Col-0) genomic DNA as template. The PCR product, which contains a 988 bp of the promoter region upstream of the 5′UTR of the *MGE2* gene, was cloned into pCR8/GW/TOPO (Invitrogen) to yield pCR8-AtMge2. To generate *MGE2* without the exonized intron within exon 1, two PCR fragments were generated with two pairs of gene-specific primers, AtMge2 5′P-1F plus AtMge2D2R and AtMge2D2F plus AtMge2 3′P-1R (Supplemental Table I) using pCR8-AtMge2 as template and cloned into pCR8/GW/TOPO. The two fragments were then ligated to yield pCR8-AtMge2D. The cloned *MGE2* and *MGE2Δ* genomic DNA were subcloned into pBGW,0 (Karimi et al., 2005) to yield the binary vectors pBGW-AtMge2 and pBGW-AtMge2D, respectively, which were then transferred into *Agrobacterium tumefaciens* GV3101 strain for transformation of Arabidopsis *mge2-1* (SALK_075614). Transformation, selection, and propagation of transgenic plants were performed as described previously (Charng et al. 2007). T3 seeds of the transgenic plants with homozygous single T-DNA insertion event were used for analysis.

### Immunoblotting

The methods of protein extraction and immunoblotting were performed as described previously (Chi et al. 2009). Antibodies against Arabidopsis MGE2 and tubulin were described previously (Hu et al. 2012; Chi et al. 2009).

### Thermotolerance assay

For TMHT assay, sterilized Arabidopsis seeds were sown on 0.5 × Murashige and Skoog medium plates containing 0.1% sucrose and imbibed at 4 °C for 3 d in the dark. The seeds were grown at 22 °C with 16 h/8 h light/dark cycle (120 μmol m^−2^ s^−1^) for 5 d before transferring to a growth chamber with temperature set at 35±0.3 °C under the 16 h light cycle and 33.5±0.3°C during the 8-h darkness. After 9 d of the prolonged heat stress treatment, the seedlings were transferred back to the previous growth condition at 22 °C and recovered for up to 10 d before phenotype was assessed.

## ACKNOWLEDGEMENTS

We thank the Transgenic Plant Core lab, Academia Sinica, for Arabidopsis transformation. Y.Y.C. acknowledges support from the intramural funding of Academia Sinica.

---

**Supplemental Fig. 1 Comparison of thermotolerance between the *E. coli grpE* mutant expressing MGE proteins with or without *Strep*-tag.** Heat-sensitive *E. coli* DA16 cells transformed with constructs encoding MGE1 and MGE2 were denoted as 1 and 2, respectively. V, DA16 transformed with empty vector. Cell cultures were serially diluted by 100-fold and spotted onto LB plates for overnight incubation at 30 °C or 43 °C.

**Supplemental Fig. 2 Purification of recombinant Arabidopsis MGE proteins.** Recombinant MGE proteins fused to *Strep*-tag were overexpressed in *E. coli* BL21 (DE3) and sequentially purified by affinity (AF), ion exchange (not shown), and gel filtration (GF) chromatographies. The purity of the proteins were evaluated by SDS-PAGE and Coomassie blue staining.

**Supplemental Fig. 3 The thermal unfolding transition of the purified recombinant MGEs with swapped domains.** A,C,E,G,I,K, the recombinant MGE variants were heated from 10 °C to 90 °C (black square) and then cooled from 90 °C to 10 °C (white square). Changes of the α-helical structure was monitored at 222 nm and expressed in mean molar ellipticity after circular dichroism (CD) spectroscopy analysis. B,D,F,H,J,L, CD spectra measured at 10 °C (solid black square), 90 °C (empty white square) and the midpoint of thermal transition (half-black square).

