## Supplemental Table 1 for "Divergence in Thermostability of Arabidopsis Mitochondrial Nucleotide Exchange Factors Encoded by Duplicate Genes, *MGE1* and *MGE2*"

**Supplemental Table I.** Primers used in this study.

| Primer name | Sequence (5'→3') |
|---|---|
| AtMge1-bs1 | GTTGGTCTCCAATGCATGGAACTCTAGATTTGAATGTTCTTC |
| AtMge1-bs4 | GTTGGTCTCCGCGCTAGCTGCAGACTCTTTGCCGC |
| AtMge2-bs3 | GTTGGTCTCCAATGTCTATGATGGATTCGTTTGCAC |
| AtMge2-bs4 | GTTGGTCTCCGCGCTAGCATCAGACTCTTTCTTTTCTTCTTG |
| pASK3+ FP | GAGTTATTTTACCACTCCCT |
| pASK3+ RP | CGCAGTAGCGGTAAACG |
| AtMge2IntD-FP | GATTCTGAGAGTGATGATGATGAATTGTC |
| AtMge2IntD-RP | ACCTGGTTGGTTTGCCTCAGCTG |
| AtMge2 5'P-1F | GTCGACGATAGGTATCTTCTATCAAACC |
| AtMge2 3'P-1R | CATATGGTTGTCTTGTTTTGAAACTGAG |
| AtMge2D2F | CCCGGGGATTCTGAGAGTGA |
| AtMge2D2R | CCCGGGTTGGTTTGCCTCAG |
