## Supplemental figure 1-3 for "Divergence in Thermostability of Arabidopsis Mitochondrial Nucleotide Exchange Factors Encoded by Duplicate Genes, *MGE1* and *MGE2*"

### Supplemental Fig. 1

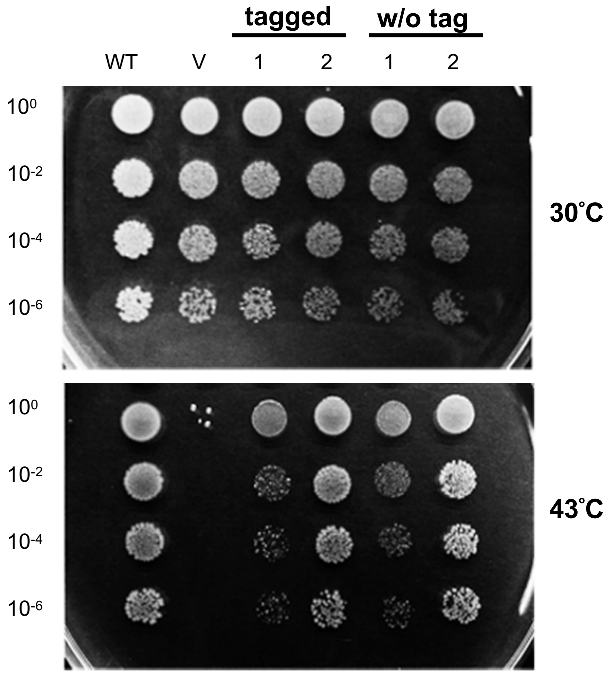


**Supplemental Fig. 1 Comparison of thermotolerance between the *E. coli grpE* mutant expressing MGE proteins with or without *Strep*-tag.**
Heat-sensitive *E. coli* DA16 cells transformed with constructs encoding MGE1 and MGE2 were denoted as 1 and 2, respectively. V, DA16 transformed with empty vector. Cell cultures were serially diluted by 100-fold and spotted onto LB plates for overnight incubation at 30 °C or 43 °C.

### Supplemental Fig. 2

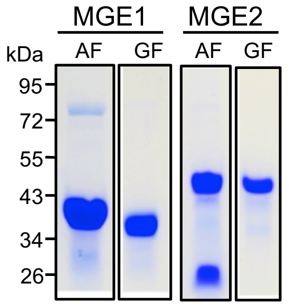


**Supplemental Fig. 2 Purification of recombinant Arabidopsis MGE proteins.** Recombinant MGE proteins fused to *Strep*-tag were overexpressed in *E. coli* BL21 (DE3) and sequentially purified by affinity (AF), ion exchange (not shown), and gel filtration (GF) chromatographies. The purity of the proteins were evaluated by SDS-PAGE and Coomassie blue staining.

### Supplemental Fig. 3

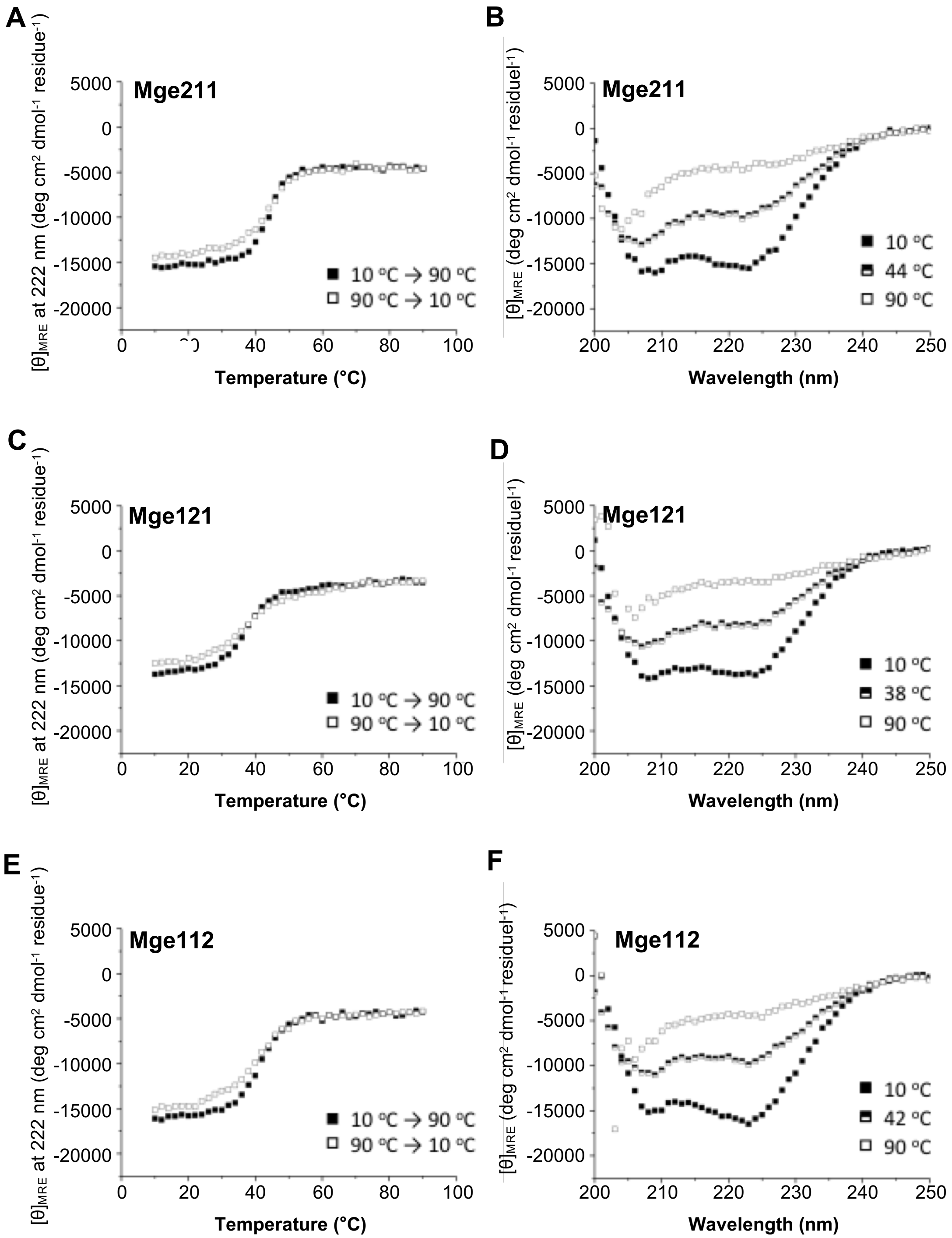


**Supplemental Fig. 3 The thermal unfolding transition of the purified recombinant MGEs with swapped domains. A,C,E,G,I,K**, the recombinant MGE variants were heated from 10 $^{o}$C to 90 $^{o}$C (black square) and then cooled from 90 $^{o}$C to 10 $^{o}$C (white square). Changes of the α-helical structure was monitored at 222 nm and expressed in mean molar ellipticity after circular dichroism (CD) spectroscopy analysis. **B,D,F,H,J,L**, CD spectra measured at 10 $^{o}$C (solid black square), 90 $^{o}$C (empty white square) and the midpoint of thermal transition (half-black square).

### Supplemental Fig. 3 (continue)

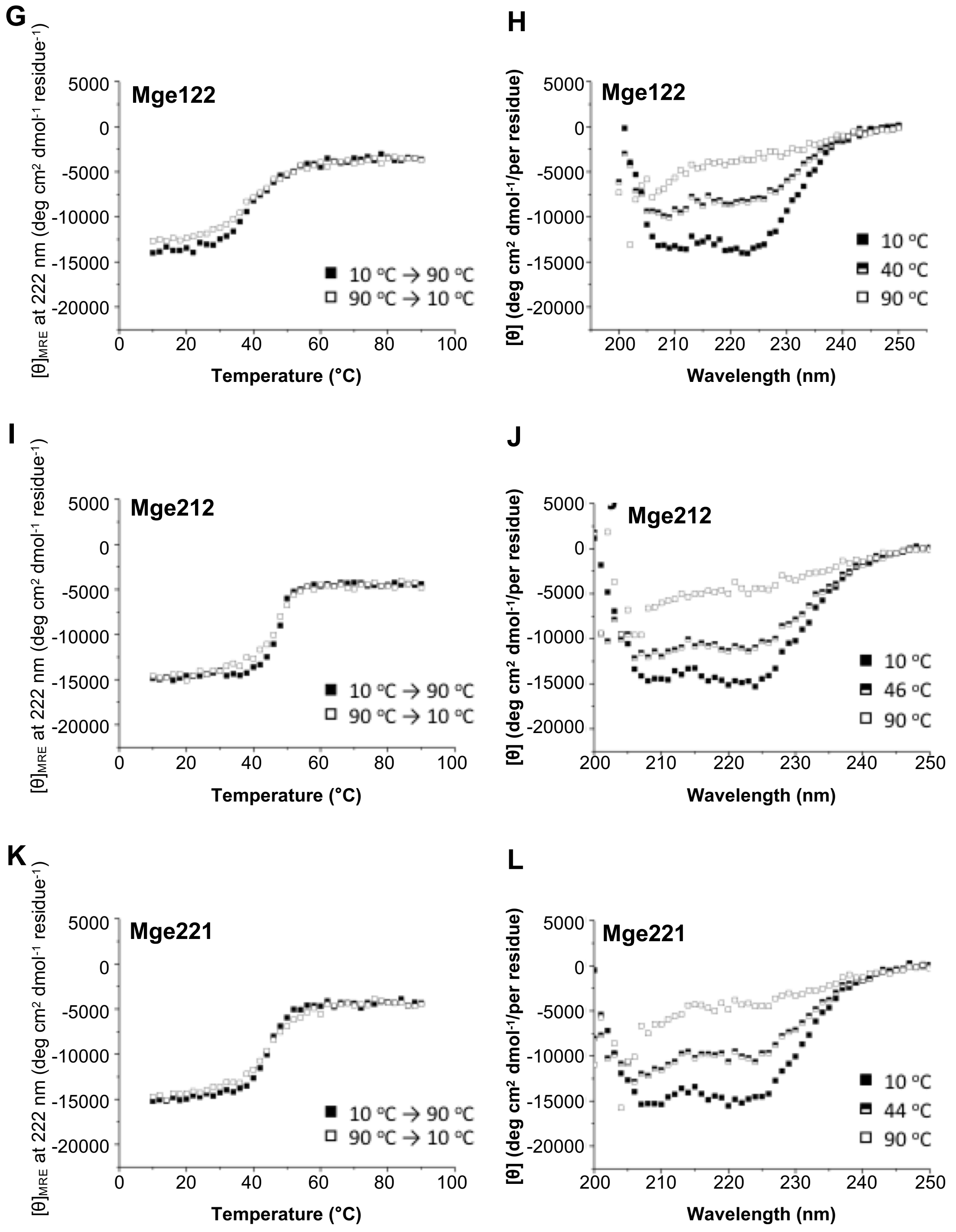
